# Neural Oscillatory Characteristics of Feedback Associated Activity in Globus Pallidus Interna

**DOI:** 10.1101/2021.12.07.471662

**Authors:** Hadi Choubdar, Mahdi Mahdavi, Zahra Rostami, Erfan Zabeh, Martin J. Gillies, Alexander L. Green, Tipu Z. Aziz, Reza Lashgari

## Abstract

Neural oscillatory activities in basal ganglia have prominent roles in cognitive processes on local and global scales. However, the characteristics of high frequency oscillatory activities during cognitive tasks have not been extensively explored in human Globus Pallidus internus (GPi). This study aimed to investigate amplitude and interhemispheric coupling of bilateral GPi high gamma bursts in dystonia and Parkinson’s Disease (PD) patients, in on and off medication states, after feedback during the Intra-Extra-Dimension shift (IED) task. Bilateral GPi Local Field Potentials (LFP) activity was recorded via externalized DBS electrodes during the IED task. Inter hemisphere phase synchrony was assessed using Inter-Site Phase Clustering (ISPC). Transient high gamma activity (∼100-150Hz) was observed immediately after feedback in the dystonia patient. Moreover, these bursts were phase synchronous between left and right GPis with an antiphase clustering of phase differences. In contrast, no synchronous high gamma activity was detected in the PD patient with or without dopamine administration. The off-med PD patient displayed enhanced low frequency clusters ameliorated by medication in the on-med state. Furthermore, an increased low frequency activity was observed after feedback of incorrect trials in both disease states. The current study provides a rare report of antiphase homotopic synchrony in human GPi, potentially related to incorporating and processing feedback information. The absence of these activities in off and on-med PD indicates the potential presence of impaired medication independent circuits related to feedback processing. Together, these findings are helpful in pointing to the potential role of GPi’s synchronized high frequency activity in cognitive tasks and feedback information processing.

## Introduction

Neural oscillatory activities are fundamental attributes of motor, behavioral, and cognitive functions of the central nervous system (CNS) (1, 2). These oscillations have been observed in various brain regions from the cortex to subcortical areas. On a local scale, oscillations are thought to be involved in the encoding and decoding of information in brain regions (3, 4). Furthermore, global scale synchronous oscillatory activity of neural circuits might play a major role in forming consciousness. Even though animal electrophysiology studies have extensively investigated the roles of neural oscillations in subcortical CNS regions, human studies are rare in this regard. Although EEG recordings are widely available, their spectral recording range is limited, especially for high-frequency oscillations. Moreover, EEG and MEG are more suitable for investigating cortical structures and have limited power for studying subcortical areas (5, 6). Recordings obtained from deep brain stimulations (DBS) have provided a precious window into investigating neural dynamics in subcortical regions (7, 8). In contrast, deep brain stimulations (DBS) electrodes offer high spatial and temporal resolution from a small target area allowing for precise studying of deep brain nuclei in real-time (9).

Parkinson’s disease is primarily caused by a reduction of dopaminergic neurons in the basal ganglia (10). Prominent symptoms in the initial stages mainly include motor dysfunctions, such as tremors and rigidity, which are superimposed by cognitive impairment and dementia as the disease progresses (10). Dystonia patients, on the other hand, suffer from sudden muscle contractions that can last from seconds to hours with mild to no apparent cognitive deficit (11). Abnormal oscillatory activities have been observed in PD and dystonia patients, especially in the GPi, and are taught to be associated with the underlying neural alterations (12). PD patients often display pathologically enhanced beta power in GPi connected to motor dysfunction severity and is temporarily alleviated by intentional motor planning and levodopa treatment (13, 14). On the other hand, Dystonia patients mainly display abnormal GPi alpha oscillations (1). DBS stimulation of GPi can alleviate motor symptoms in both dystonia and PD, especially in severe and treatment-resistant cases (10, 11).

Although GPi is mainly known for its role in motor movement and coordination, recent studies have also proposed a cognitive role for this region which points to possible functionality for cognitive processing of action and response selection, especially during high conflict states (15). With a general functional viewpoint, previous reports have demonstrated that gamma and theta band activities are closely linked with memory processes, whereas alpha and beta oscillations have been robustly observed in attention-centered tasks (16, 17). Besides, more recent studies have expanded findings regarding oscillatory CNS roles by proposing a prominent link between high gamma bursts, also known as ripples, in cortical and several subcortical regions, such as the hippocampus with memory encoding and recall. Transient high gamma activity in the hippocampus has been linked to explicit awareness of the working memory content and successful memory task performance. Many animal models (18) and several human studies have studied this mnemonic function of high gamma bursts, and their synchrony with cortical structures has been studied in many animal models (19) and several human studies (20, 21).

The cognitive role of high gamma GPi activities in humans has not been fully explored. Gillies et al. observed high gamma activity in dystonia patients after receiving audiovisual feedback during the Cambridge Cognition’s IED task (22). In this study, we aimed to delve deeper into the characteristics of high gamma oscillatory bursts in GPi during the Intra Extra Dimensional task (IED) in recordings from PD and dystonia patients. More specifically, we evaluated the amplitude and inter-hemisphere coupling characteristics of high gamma bursts between left and right GPis after receiving feedback. Furthermore, the results were compared between dystonia and PD as normal and dopaminergic deficit states, respectively.

## Materials and methods

### Patients

This study uses a subset of data previously reported by Gillies et al. (22). Two patients were included in the study (Table. 1). both patients were female and right-handed. The first patient was 59 years old with focal cervical dystonia. The second one was a 66 years old Parkinsonian patient. The dystonic patient received no medication during the test period. The parkinsonian patient first underwent the test before receiving scheduled prescribed L-dopa in order to simulate “off-med” symptoms. Subsequently, the test was repeated after scheduled prescribed L-dopa medication when the patient was experiencing “on-med” symptoms. Both patients gave informed written consent. The study was conformed to the Declaration of Helsinki and approved by Oxfordshire Research Ethics Committee A.

**Table 1.** Patients’ Characteristics

|  | Diagnosis | Gender | Surgery Age | First reported symptom age | Handedness | NART IQ | AMIPB (delayed) |
| --- | --- | --- | --- | --- | --- | --- | --- |
| 1 | Focal Dystonia (cervical) | Female | 59 | 35 | Right | n/a | n/a |
| 2 | Parkinson's Disease (On and off-med) | Female | 55 | 47 | Right | 103 | Pass (9) |

All scores have a mean 100 and SD 15 unless otherwise stated. Performance is classified as: impaired (<69), borderline (70–79), low average (80–89), average (90–109), high average (110–119), superior (120–129), very superior (>130).
NART IQ National adult reading test intelligence quotient, AMIPB adult memory and information processing battery.

### Surgical Procedure

Both patients underwent bilateral Globus pallidus deep brain stimulation via a standard technique (23). Medtronic 3387 ® DBS leads were placed in left and right posteroventral GPi. Each electrode had four circumferential 1.5 mm channels. A CT head scan was performed to assess the lead position after implantation (verified by Image Fusion with the pre-operative MRI). Internalization of DBS leads and implantation of internal pulse generators was done approximately a week after clinical testing for efficacy (fig. 1A).

**Figure 1:**
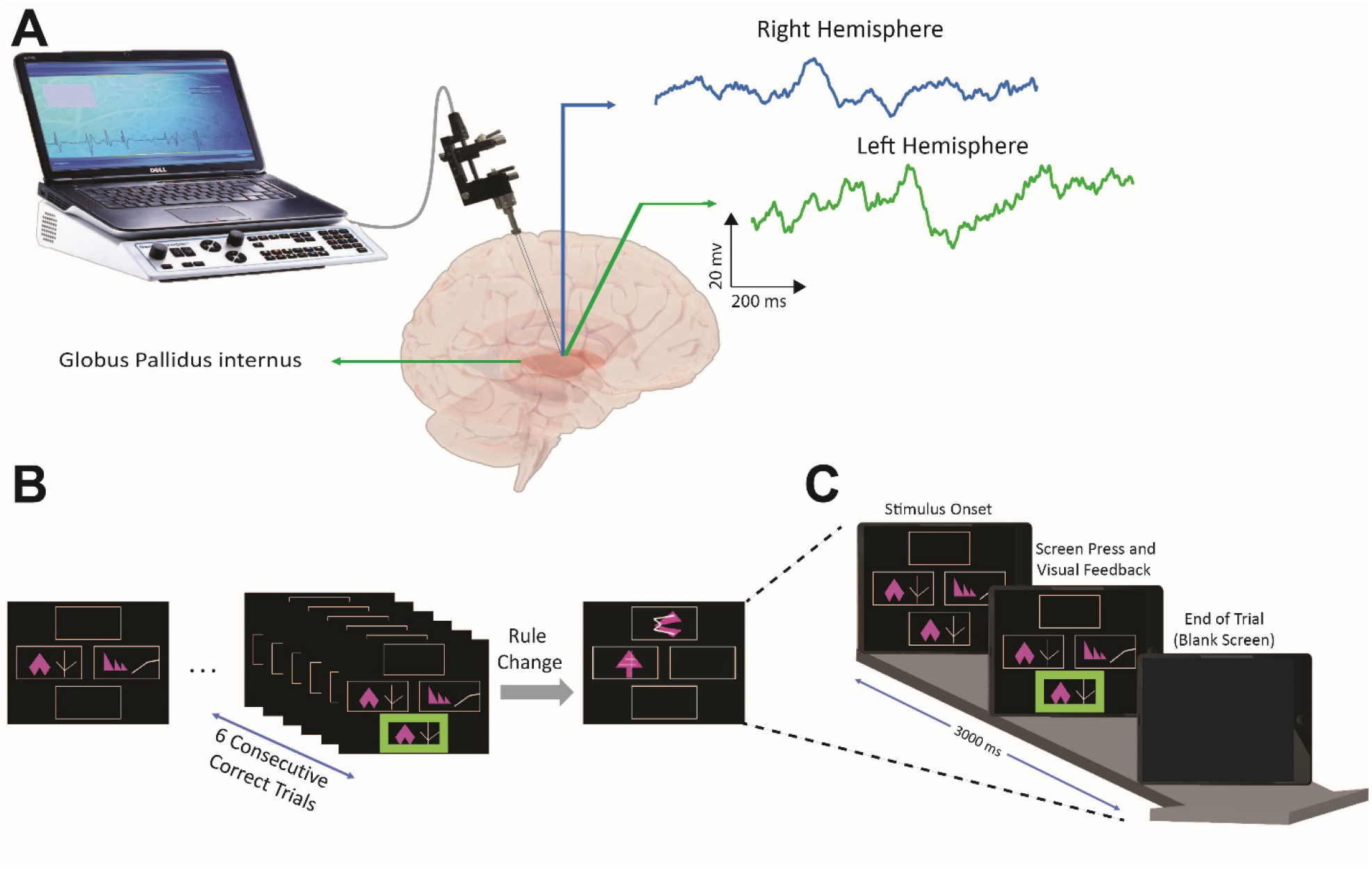
The overview of data recording and IED task. **A)** two DBS leads with three recording channels were placed in the right and left GPi separately to record the LFP signal during the IED task. **B)** schematic of Cantab^®^ Intra-Extra Dimension shift task. The task contains nine rule sets. In each section, when the patient completes six correct trials sequentially, the rule changes. Each rule section has 50 trials at maximum, and if the patient does not reach six consecutive correct trials within the 50 trials, the experiment ends at that rule section. **C)** Graphic representation of a single trial. Each trial starts when the visual object appears on the blank screen. The patient moves to touch the screen in order to choose the preferred object. Auditory and visual feedbacks are given immediately and simultaneously to the patient upon screen touch. The red marker indicates the “incorrect” answer in the visual feedback, and the green marker indicates the “correct” answer. For auditory feedback, the patient hears “correct” for the correct answers and “incorrect” for incorrect answers. The trial ends when the screen becomes blank before the start of the next trial.

### Experimental Procedure

Patients executed a forced decision-making task, Intra-Extra Dimension shift task (by Cambridge Cognition), an on-screen variation of the Wisconsin card sorting task. The dystonic patient performed the task once, and the parkinsonian patient performed the task in both on-med and off-med conditions. The IED task contains nine rule sections, changing intra-dimensionally until the 7^th^ section and changing extra-dimensionally in the 8^th^ and 9^th^ sections. Within each section, the subject experience several trials up to a maximum of 50 trials. The six consecutive correct answers to trials lead the subject to pass the section and change the rule. If the subjects do not achieve six subsequent correct answers within 50 tries, they fail the section, and the experiment ends (fig. 1B). Each trial starts when a visual object containing two abstract figures appears on the blank screen. The subject moves his hand to touch the screen and choose the answer (fig. 1C). The subject receives simultaneous audiovisual feedback immediately after the answer selection. The visual feedback presents as green and red highlights on the screen for “correct” and “incorrect” answers, respectively. The subject hears “correct” for “correct” answers and “wrong” for “incorrect” answers as auditory feedback. After receiving the feedbacks, the trial ends when the screen becomes blank. Each trial lasted about 3000 to 3500ms with 500ms of blank screen presentation.

### Electrophysiological recording

Recordings were obtained from adjacent circumferential 1.5 mm contacts of each deep brain macroelectrode in a bipolar configuration to reduce the effects of volume conduction (9). Globus pallidus contacts were identified by postoperative image-fused MRI and CT. Signals were amplified (10,000×) using isolated CED 1902 amplifiers and digitized via CED 1401 Mark II at a rate of 2.5 kHz (Cambridge Electronic Design) or recorded by a Porti system (Twente Medical Systems International, BV, Netherlands) and recorded onto storage using Spike2 software (Cambridge Electronic Designs, Cambridge, UK). Raw data was notch filtered at 50, 100, and 150 Hz as required using Spike2 infinite impulse response Bessel filters, *Q* value adjusted to avoid unwanted filtering of adjacent frequencies as much as possible. Spike 2 data were imported into EEGLAB (24).

### Data Analysis

All analysis steps were performed using MATLAB R2019b. To inspect trials’ signal quality, image matrices of trials were visually inspected. Consequently, one trial from the dystonic patient and three trials from the on-med PD patient were removed. A high pass filter at 0.1Hz and a low pass filter at 200Hz were also utilized to remove slow and fast oscillatory noises. The first channel of each electrode was used for referencing. To perform time-frequency decomposition, Morlet wavelets with increasing cycles were utilized over 120 frequency steps between 2 and 150 HZ. The baseline window was designated to be 2200-2500 ms after the start of each trial during the blank screen phase. Amplitude values were converted to decibels using the mean amplitude calculated in the aforementioned window for each trial.

To find statistically significant time-frequency activities, cluster correction with a time-frequency-amplitude voxel and cluster threshold of 0.05 was implemented. Briefly, distribution for each voxel was obtained in 1000 iterations. In each iteration, the amplitude spectrum of each trial was cut in a random time point, the order was swapped, and the mean amplitude was calculated over trials. After the initial permutation loop, original amplitude values were converted into z-scores using the mean and standard deviation of the acquired distribution. In the next step, to obtain a distribution for amplitude cluster sizes, another 1000 iteration was implemented. In each iteration, values from the previous loop were converted to z score, and a pixel threshold of 0.05 was utilized, after which the size of the largest cluster was stored. After the permutation loop, only clusters larger than the 95^th^ percentile were considered significant.

To obtain the standard error of the mean (SEM) for each band, bootstrapping over trials in 1000 iterations was used. In brief, in each iteration, an equal number of trials as the original state were selected with replacement, and the SEM was calculated over trials. Afterward, all SEM values were averaged across iterations to obtain the final SEM. To identify time points where the difference between the amplitude time series of two patients was significantly different, the normalized amplitude time series were subtracted, and bootstrapping over 1000 iterations was used to obtain confidence intervals (CIs). Afterward, windows, where the subtracted time series and the CI did not envelope 0, were designated as statistically significant.

To investigate the phase synchrony between left and right GPis, Inter-Site Phase Clustering (ISPC) was implemented. Phase differences were calculated from the wavelet convolution and were converted to complex representation using the formula 1. The calculated representations were averaged in sliding time windows varying according to the frequency cycle in each trial, and the results were averaged over all trials with the same feedback status. This approach is robust to trial to trial jitters and can measure the total (phase-locked and none phase-locked) connectivity. The resulting values were differentiated from the mean ISPC values from the baseline window to remove baseline effects. Permutation testing with a threshold of 0.05 in a manner similar to the abovementioned method was used for statistical testing (25).

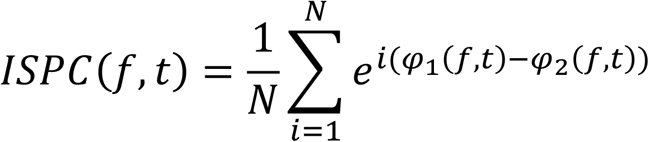

## Results

### Frequency band characteristics of oscillatory activities

Time-frequency analysis demonstrated a transient activity in the high gamma band (60-140 Hz) immediately after the feedback. Cluster correction on the time-frequency plot showed the significance of the high gamma oscillation (p-value<0.05) (fig. 2A). However, there was no significant high gamma activity in the Parkinsonian patient in the equivalent time window (fig. 2B). Furthermore, the amplitude comparison between correct and incorrect trials with respect to high gamma oscillation in the dystonic patient showed no significant difference between correct and incorrect trials (p-value>0.05), indicating the robustness of the high gamma activity to the received feedbacks (fig. 2C).

**Figure 2:**
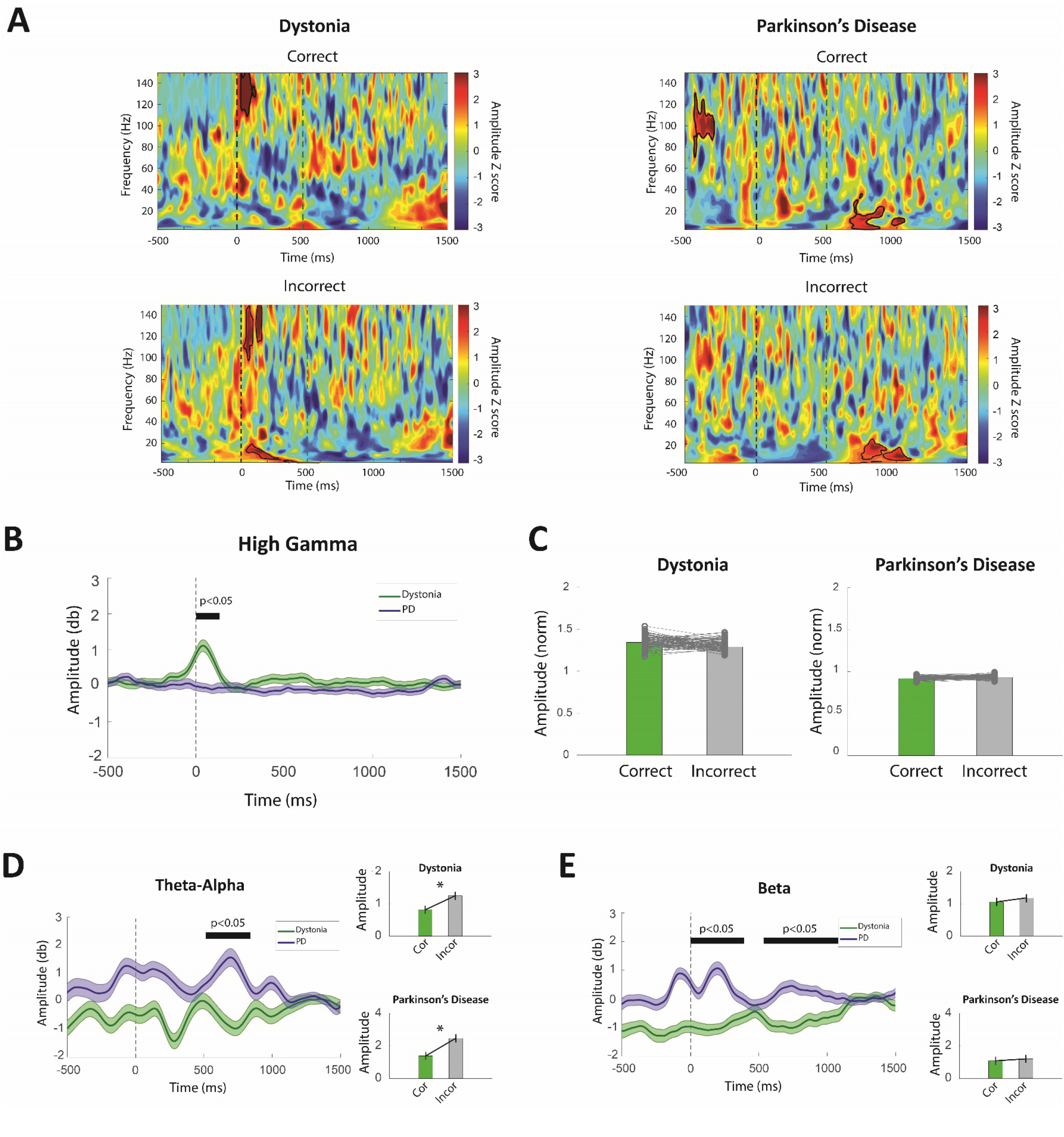
Activity dynamics of frequency bands in dystonia and Parkinson’s disease during the IED task. **A)** Cluster corrected time-frequency amplitude spectrum of dystonia (left) and PD (right) patients during correct and incorrect trials. Immediately after receiving feedback, a significant cluster of high gamma activity (permutation test; P < 0.05) is observed in the dystonia patient during correct and incorrect trials. In contrast, no significant cluster was observed in the same time-frequency window in the PD patient. A significant cluster is observable in the PD patient around 600 ms in the theta-alpha and beta frequency bands, most likely due to the pathophysiology of the disease. **B)** Average high gamma amplitude time series for the dystonia and PD patient shows that high gamma burst is significant in the dystonia patient (permutation test; P < 0.05), marked by the black bar above the plot. **C)** Average high gamma amplitudes were calculated for the correct and incorrect trials in the [0, 200] ms time window. Bar plots of average amplitudes showed no difference between correct and incorrect trials in both dystonia and the PD patient. **D)** Averaged theta-alpha amplitude in the dystonia and PD patients. The PD patients displayed enhanced low frequency activity in time windows after the feedback (permutation test; P < 0.05) marked by the black bar above the plot. Furthermore, a comparison of average amplitudes in correct and incorrect trials ([0, 200]ms for dystonia and [500, 1000]ms for the PD patient) revealed a significantly higher theta-alpha activity in both patients during the incorrect trials (permutation test; P < 0.05). **E)** Averaged beta amplitude time series in the dystonia and PD patients. The PD patients displayed enhanced beta frequency activity after the feedback (permutation test; P < 0.05) marked by the black bar above the plot. No difference between correct and incorrect trials ([500 1000]ms for both patients) in both dystonia and PD patients was seen. To avoid potential bias effects related to the treatment medication, only the activity from the off-med PD patient was utilized in this figure.

Furthermore, a low frequency activity in the theta, alpha, and beta frequency band (∼ 4-30 Hz) was detected in the PD patient’s time-frequency analysis. Further cluster correction demonstrated the significance of this activity (p-value<0.05). Compared to the dystonic patient, the current activity in low frequency bands is significantly higher in the PD patient (p-value<0.05) (fig. 2D, E). Moreover, alpha-theta activity in both Dystonic and PD patients was significantly higher in incorrect trials than in correct trials (p-value<0.05) (fig. 2D). However, there was no significant difference in beta frequency activity between correct and incorrect trials in those patients (p-value<0.05) (fig. 2E).

### Inter-hemisphere synchrony in high gamma oscillation

Inter-site phase clustering (ISPC) was used for evaluating phase synchrony between right and left GPi (fig. 3A). The results of the dystonic patient showed that there is a statistically significant (p-value<0.05) phase synchrony between left and right GPi in the high gamma frequency band immediately after feedback (fig. 3B). In contrast, there was no significant phase synchrony between right and left GPi in the PD patient’s high and low frequency bands (fig. 3B).

**Figure 3:**
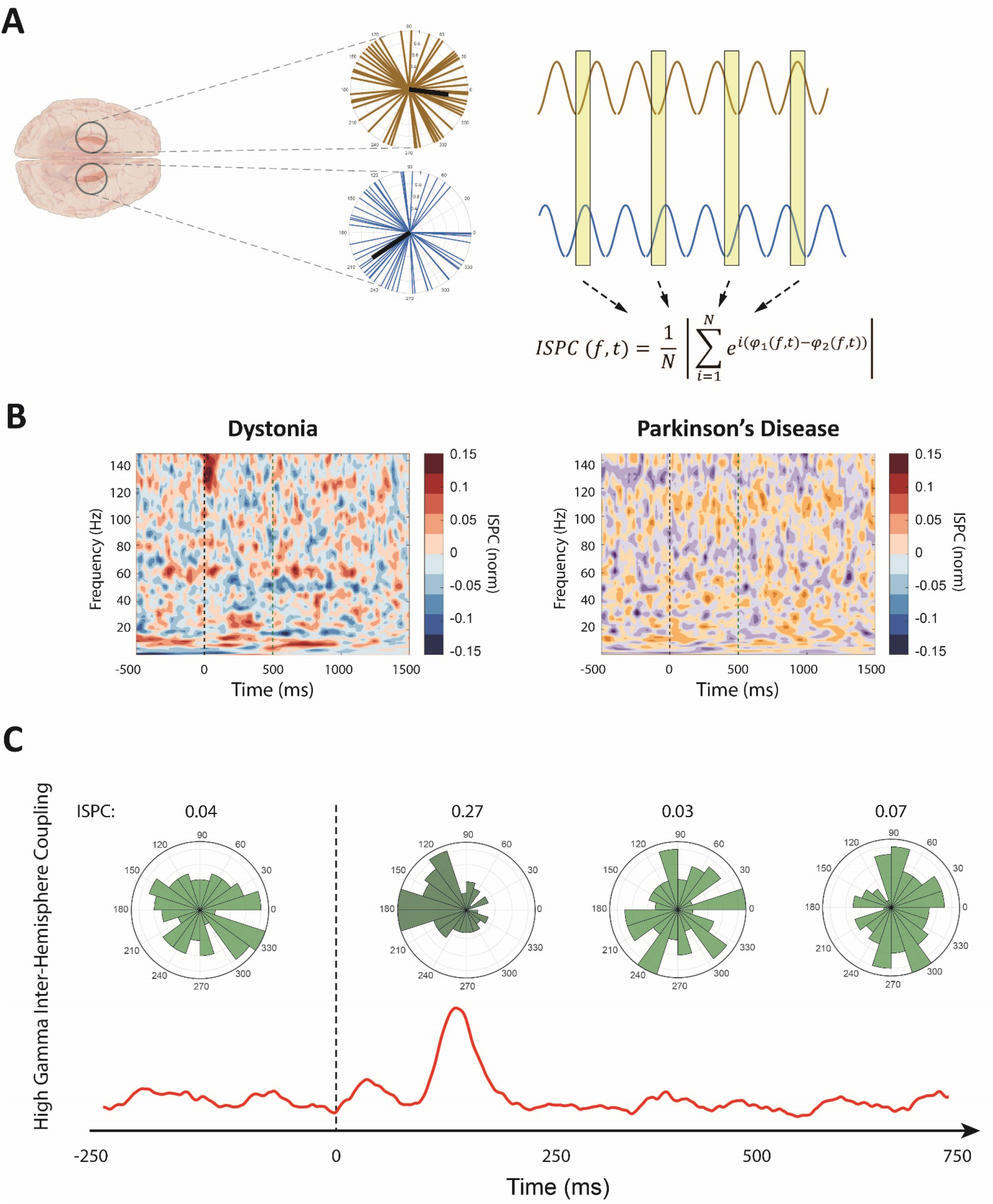
Phase synchrony between left and right GPi in dystonia and PD patients during the IED task. **A)** schematic overview of Inter-Site Phase Clustering (ISPC) method for calculating phase synchrony. At each time-frequency point, the phase difference between two sites is transformed via Euler’s formula and is averaged over trials. **B)** ISPC synchrony plots of dystonia and the PD patient. Immediately upon feedback, prominent synchrony between right and left GPi in the high gamma frequency is observable in the dystonia patient but not in the PD patient. **C)** Polar histogram of the phase difference between left and right GPi in high gamma frequency band in the dystonia patient. A conspicuous clustering of phase differences around 180° is present immediately after the feedback. In the same time window, the average high gamma activity shows a marked peak. To avoid potential bias effects related to the treatment medication, only the activity from the off-med PD patient was utilized in this figure.

While ISPC can detect the presence of phase synchrony between two sites, it does not reveal the clustering of the phase differences between the two measurement areas. Therefore, in order to investigate the distribution of phase differences between left and right GPi, the phase differences in each trial were converted to Euler representations and were subtracted by the average representation in the trial’s baseline window to remove the baseline effects. Then the representations were averaged over the high gamma spectrum, and average phase difference angles were extracted from the resulting representations. Finally, the angles were binned into a polar histogram to show the distribution of average phase differences between right and left GPi. The results indicated that the observed high gamma phase synchrony in the dystonia patient exhibited an antiphase clustering of bilateral phase differences in the proximity of 180° (fig. 3C). The average ISPC in the high gamma band was also consistent with this result (fig. 3C).

### Medication effect on the low frequency oscillation in PD patient

The PD patient showed different characteristics in low frequency oscillations in the on-med and off-med states. Results of the cluster corrected time-frequency analysis demonstrated that in contrast to the off-med state, low frequency activity in theta, alpha, and beta frequency bands is not significant in the on-med state (fig. 4A). In order to further investigate, a permutation test was performed between on and off-med states in each frequency band. There was a significant difference between on and off-med states in the alpha-theta band (p-value<0.05). The same result was repeated in the beta frequency band after the feedback (p-value<0.05). However, there was no significant difference in the high gamma frequency band between on and off-medication states (fig. 4B).

**Figure 4:**
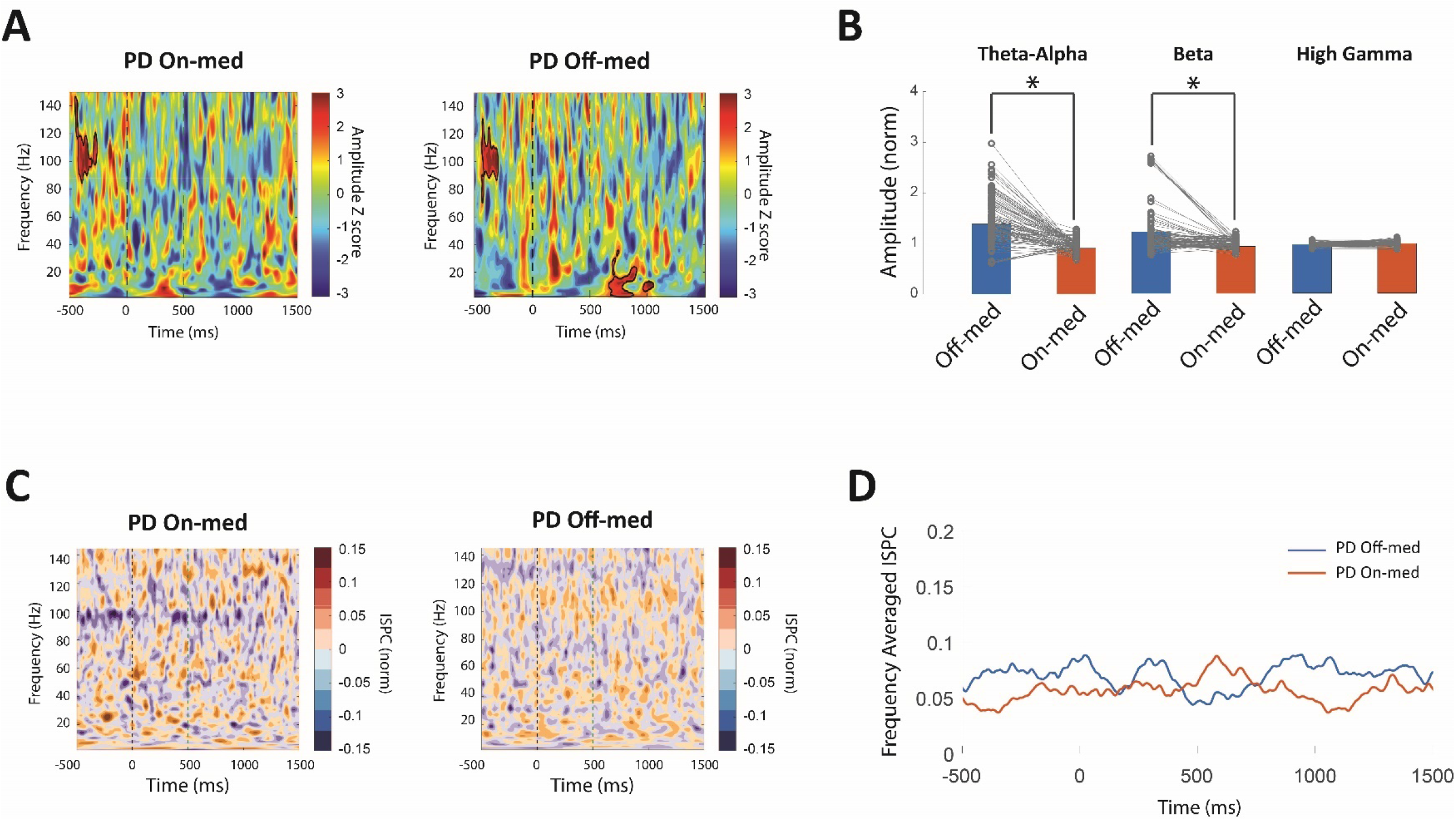
Medication effect on the activity dynamics and phase synchrony of frequency bands in on-med and off-med PD patients during the IED task. **A)** The observed alpha and beta activity cluster in the off-med patient is absent in the on-med patient, which is in accordance with previous reports indicating the effect of medication on alleviating pathological low frequency bursts in the PD patients. No high gamma cluster was observed after the feedback in both patients. **B)** Using the panel A results, average amplitudes were calculated in the [500, 1000] ms time windows for the theta-alpha and beta bands and the [0, 500]ms window for the high gamma band. The results revealed a significant difference of average amplitude between on-med and of-med states (permutation test; P<0.05) but no difference in the high gamma activity. **C, D)** ISPC synchrony time-frequency spectrum and frequency averaged high gamma ISPC time series of the PD patient in on and off med states displayed no alteration in the phase synchrony state between right and left GPi with medication in PD patient.

Furthermore, ISPC analysis was performed to evaluate the inter-hemisphere phase synchrony and compare them between two medication states. Similar to the off-med state, there was no significant high gamma phase synchrony between right and left GPi in the on-med state of the PD patient in both ISPC time-frequency spectrum (fig. 4C) and average high gamma ISPC time series (fig. 4D).

## Discussion

This study investigated the characteristics of LFP recordings from bilateral GPi in dystonia and Parkinson’s disease during the performance of the IED task. Exploring the roles of deep brain structures in humans during a cognitive task is an arduous process. DBS data provide a valuable window for direct investigation of basal ganglia but are rare due to the small number of DBS surgeries and the effort required on behalf of patients and researchers for proper recording and avoiding complications. Therefore, observations from DBS recordings are valuable sources of information for investigating the role of subcortical areas in cognitive functions. A noticeable finding of this study was the presence of short, high gamma activity bursts after receiving audiovisual feedback in the dystonia patient. We further evaluated this finding by using a robust cluster detection statistical test to ensure correction for multiple comparisons. The observed activity was not affected by the semantic nature of the feedback as both correct and incorrect feedbacks elicited the phenomena. The same observation, however, was not detected in the PD patient, both in on and off medication states. These results are similar to the findings of Gillies et al. (22), where similar patterns were observed. Furthermore, an interesting finding was that the observed high gamma activity showed marked phase synchrony between bilateral GPis immediately after the feedback in the dystonia patient during both correct and incorrect trials. The observed synchrony in the dystonia patient had an antiphase characteristic with a prominent binning of phase differences around 180°. In contrast, this synchrony was absent in the PD patient in both on-med and off-med states. Recent studies have reported the presence of antiphase synchrony in homotopic cortical structures. However, to the best of our knowledge, the current study provides the first investigation of homotopic antiphase synchrony in human GPi in dopamine-intact and dopamine-deficit states.

Although GPi is mainly known as a motor center, previous studies have reported that it is also involved in non-motor cognitive tasks, such as working memory and attention, besides its motor functionality (15). High gamma oscillations are often associated with increased activity of local neural populations (26, 27). Previous results have mainly linked high gamma activity with various aspects of the memory. For example, Animal studies in rodents have demonstrated high gamma activity in several cortical and subcortical regions that were mainly linked with long-term memory performance (19). Similar high gamma activity patterns have also been observed in studies on humans. A study by Norman et al. using iEEG recording on human patients during the execution of a memorization-recall task observed a brief high gamma ripple activity, also termed Sharp Wave Ripples (SWR), in the hippocampus (28). The subjects were asked to observe a number of pictures and recall them after a period of time. The results revealed that high gamma SWRs are key players in the encoding memory process, and the ability of a patient to remember a picture successfully was linked to the high gamma burst activity during the recall phase. Furthermore, the observed bursts displayed synchronous activity with several cortical regions, such as early visual areas and fusiform gyrus. The authors concluded that high gamma ripples create a dialogue between the hippocampus, a subcortical area, and cortical regions involved in memory processing. A cornerstone of the IED task is the ability to successfully recall the results of recent audiovisual feedback, which is directly related to the subject’s working memory (29). As a result, it is likely that the observed high gamma activity of the current study, at least in part, is related to the process of memory formation and recalling of the recently received feedback in order to encode the information to be used in the upcoming trial.

One of the main findings of this study that adds to the previous literature was the presence of antiphase synchrony in the observed high gamma burst between right and left GPi in the dystonia patient. Recent studies have reported prominent antiphase synchrony in intracranial EEG recordings between homotopic regions (30, 31). Antiphase synchrony may be involved in augmenting the brain’s predictive capability and task performance improvement by synchronizing the activity of neural circuits in response to internal or external cues. Furthermore, the observed functional connectivity could serve as an integration mechanism across separate brain regions (30). The observation that these brief synchrony epochs rapidly appear after feedback and are present in trials regardless of the semantic nature of the received feedback is supportive of the integrative role of the observed connectivity. The aforementioned processes are similar to the back-propagation of an artificial neural network (ANN), where the loss is computed in the last layer and then is back-propagated across the networks to optimize the framework by adjusting the units of each layer simultaneously (32). The higher cortical areas can be seen as the last layer of an ANN where output is calculated and compared with the desired output. To optimize the output’s precision, an intrinsic loss is calculated and propagated towards downstream centers to optimize their activity. Thus, the observed synchronous high gamma bursts can be attributed to the simultaneous arrival of optimization signals from higher cortical levels.

In contrast to the dystonia patient, no significant phase synchrony was observed between left and right GPis in the PD patient in both on-med and off-med states. As a result, it can be assumed that the high gamma synchrony system requires a structurally intact dopaminergic pathway since even treatment with dopaminergic drugs was not able to restore this synchrony. Moreover, previous studies have reported the presence of abnormal structural changes in PD patients where the severity of these changes is associated with the degree of memory dysfunction in these patients (33). Moreover, it has been suggested that an intact corpus callosum is required for the presence of antiphase synchrony between homotopic regions (30). PD patients, however, often suffer from various degrees of corpus callosum atrophy (34), which might be a potential cause for the observed lack of antiphase synchrony in bilateral GPi of these patients. From a cellular perspective, high frequency bursts have been reported to be associated with gap junction structures, especially axonal complexes. Previous studies in rodents (35) and humans (36), especially in the hippocampus, have reported the presence of gap junctions during gamma and high gamma activity. Furthermore, it is known that compensatory changes in membrane channels and junctions are present in PD patients. These changes might be associated with the observed absence of high gamma activity in the PD state, but more studies in this regard are needed as the role of gap junctions in human basal ganglia oscillatory dynamics has been barely investigated (37).

This study also observed enhanced low frequency oscillations, mainly in theta-alpha and beta frequency bands, in the off-med PD patient compared with the dystonia patient. Previous studies have reported abnormally increased low frequency activities in PD patients (13, 38). Prominent theta and alpha oscillations in the basal ganglia have been observed in association with disrupted risk assessment errors. It is possible that the observed higher theta-alpha power after the feedback during the incorrect trials in off-med PD and dystonia patients is related to reward prediction error. Another observed aspect of the PD disease is the beta oscillations that are strongly linked to motor symptoms; these oscillations are decreased during voluntary movements but rebound after the movement’s conclusion (14). Accordingly, the observed abnormal beta activity in the PD patient is likely to be related to the aforementioned beta rebound after finishing voluntary movement since it appears during the time when the subject has likely withdrawn their hand. All the abnormal low frequency activities faded during the on-med phase, pointing to the balancing effects of medication in PD disease.

The findings of this study should be viewed in the light of several limitations. The number of subjects in this study was not large. This limited the possibility of additional analyses, such as an in-depth investigation of the detailed oscillatory characteristics of different stages of the IED task. During the recording phase, only markers for the feedback time were recorded. As a result, trials were centered with respect to the feedback time. Therefore, we cannot make precise assumptions regarding the oscillatory events triggered by the stimuli presentation. The IED task might not be completely orthogonal as several cognitive factors can affect the subject’s performance during the trials. Finally, dystonia was considered a control state for the PD group since they have different underlying pathophysiology. However, it is possible that dystonia patients also suffer from a mild degree of functional impairment. Future works should evaluate the results of this study with larger study populations and use more specific tasks with respect to the study’s framework.

## Conclusion

The present study investigated oscillatory neural dynamics of GPi in dystonia and PD patients during the IED task. The results revealed the presence of bilateral antiphase synchrony in the high gamma band immediately after the feedback in the dystonia patient. No similar observation was present in the PD patient in both on-med and off-med states. High gamma activity has been observed in association with memory processing and recall during tasks. Antiphase functional connectivity between homotopic regions has been suggested to be involved in input integration and processing across distinct areas to augment the anticipatory performance of the brain. Together, the observed antiphase high gamma synchrony could be related to the feedback integration and memory processing to tune the subject’s task across trials. The absence of antiphase high gamma activity in the PD patient in both on and off med states can be associated with the dopamine administration independent structural deficits that are routinely present in disorder. Future studies are required to verify this framework using larger study populations and more specific cognitive tasks.

## References

1. Eisinger RS, Cernera S, Gittis A, Gunduz A, Okun MS. A review of basal ganglia circuits and physiology: application to deep brain stimulation. Parkinsonism & related disorders. 2019;59:9–20.

2. Halje P, Brys I, Mariman JJ, da Cunha C, Fuentes R, Petersson P. Oscillations in cortico-basal ganglia circuits: Implications for Parkinson’s disease and other neurologic and psychiatric conditions. Journal of neurophysiology. 2019;122(1):203–31.

3. Jacobs J, Hwang G, Curran T, Kahana MJ. EEG oscillations and recognition memory: theta correlates of memory retrieval and decision making. Neuroimage. 2006;32(2):978–87.

4. Flight MH. Oscillations help to decode spike patterns. Nature Reviews Neuroscience. 2009;10(12):835-.

5. Rad PN, Behzadi F, Yazdanfar A, Ghamari H, Zabeh E, Lashgari R. Cognitive and perceptual influences of architectural and urban environments with an emphasis on the experimental procedures and techniques. 2021.

6. Buzsáki G, Anastassiou CA, Koch C. The origin of extracellular fields and currents—EEG, ECoG, LFP and spikes. Nature reviews neuroscience. 2012;13(6):407–20.

7. Singh A, Kammermeier S, Plate A, Mehrkens JH, Ilmberger J, Bötzel K. Pattern of local field potential activity in the globus pallidus internum of dystonic patients during walking on a treadmill. Experimental neurology. 2011;232(2):162–7.

8. Singh A, Bötzel K. Globus pallidus internus oscillatory activity is related to movement speed. European Journal of Neuroscience. 2013;38(11):3644–9.

9. Lempka SF, McIntyre CC. Theoretical analysis of the local field potential in deep brain stimulation applications. PloS one. 2013;8(3):e59839.

10. Dauer W, Przedborski S. Parkinson’s disease: mechanisms and models. Neuron. 2003;39(6):889–909.

11. Tarsy D, Simon DK. Dystonia. New England Journal of Medicine. 2006;355(8):818–29.

12. Silberstein P, KuÈhn AA, Kupsch A, Trottenberg T, Krauss JK, WoÈhrle JC, et al. Patterning of globus pallidus local field potentials differs between Parkinson’s disease and dystonia. Brain. 2003;126(12):2597–608.

13. Yugeta A, Hutchison WD, Hamani C, Saha U, Lozano AM, Hodaie M, et al. Modulation of Beta oscillations in the subthalamic nucleus with prosaccades and antisaccades in Parkinson’s disease. Journal of Neuroscience. 2013;33(16):6895–904.

14. Engel AK, Fries P. Beta-band oscillations—signalling the status quo? Current opinion in neurobiology. 2010;20(2):156–65.

15. Navid MS, Kammermeier S, Niazi IK, Sharma VD, Vuong SM, Greenlee JD, et al. No difference in cognitive task-related oscillations between human internal globus pallidus and subthalamic nucleus. medRxiv. 2021.

16. Ward LM. Synchronous neural oscillations and cognitive processes. Trends in cognitive sciences. 2003;7(12):553–9.

17. Taghizadeh B, Foley NC, Karimimehr S, Cohanpour M, Semework M, Sheth SA, et al. Reward uncertainty asymmetrically affects information transmission within the monkey fronto-parietal network. Communications biology. 2020;3(1):1–11.

18. Fernández-Ruiz A, Oliva A, de Oliveira EF, Rocha-Almeida F, Tingley D, Buzsáki G. Long-duration hippocampal sharp wave ripples improve memory. Science. 2019;364(6445):1082–6.

19. Tingley D, Buzsáki G. Routing of hippocampal ripples to subcortical structures via the lateral septum. Neuron. 2020;105(1):138-49. e5.

20. Weiss SA, Song I, Leng M, Pastore T, Slezak D, Waldman Z, et al. Ripples have distinct spectral properties and phase-amplitude coupling with slow waves, but indistinct unit firing, in human epileptogenic hippocampus. Frontiers in neurology. 2020;11:174.

21. Pail M, Cimbálník J, Roman R, Daniel P, Shaw DJ, Chrastina J, et al. High frequency oscillations in epileptic and non-epileptic human hippocampus during a cognitive task. Scientific reports. 2020;10(1):1–12.

22. Gillies M, Hyam J, Weiss A, Antoniades C, Bogacz R, Fitzgerald J, et al. The cognitive role of the globus pallidus interna; insights from disease states. Experimental brain research. 2017;235(5):1455.

23. Yianni J, Green AL, Aziz TZ. Surgical treatment of dystonia. International review of neurobiology. 2011;98:573–89.

24. Delorme A, Makeig S. EEGLAB: an open source toolbox for analysis of single-trial EEG dynamics including independent component analysis. Journal of neuroscience methods. 2004;134(1):9–21.

25. Cohen MX. Effects of time lag and frequency matching on phase-based connectivity. Journal of neuroscience methods. 2015;250:137–46.

26. Parvizi J, Kastner S. Promises and limitations of human intracranial electroencephalography. Nature neuroscience. 2018;21(4):474–83.

27. Lashgari R, Li X, Chen Y, Kremkow J, Bereshpolova Y, Swadlow HA, et al. Response properties of local field potentials and neighboring single neurons in awake primary visual cortex. Journal of Neuroscience. 2012;32(33):11396–413.

28. Norman Y, Yeagle EM, Khuvis S, Harel M, Mehta AD, Malach R. Hippocampal sharp-wave ripples linked to visual episodic recollection in humans. Science. 2019;365(6454).

29. Oh A, Vidal J, Taylor MJ, Pang EW. Neuromagnetic correlates of intra-and extra-dimensional set-shifting. Brain and cognition. 2014;86:90–7.

30. O’Reilly C, Elsabbagh M. Intracranial recordings reveal ubiquitous in-phase and in-antiphase functional connectivity between homotopic brain regions in humans. Journal of Neuroscience Research. 2021;99(3):887–97.

31. Merrison-Hort RJ, Borisyuk R. The emergence of two anti-phase oscillatory neural populations in a computational model of the Parkinsonian globus pallidus. Frontiers in computational neuroscience. 2013;7:173.

32. Sibi P, Jones SA, Siddarth P. Analysis of different activation functions using back propagation neural networks. Journal of theoretical and applied information technology. 2013;47(3):1264–8.

33. Possin KL, Filoteo JV, Song DD, Salmon DP. Spatial and object working memory deficits in Parkinson’s disease are due to impairment in different underlying processes. Neuropsychology. 2008;22(5):585.

34. Goldman JG, Bledsoe IO, Merkitch D, Dinh V, Bernard B, Stebbins GT. Corpus callosal atrophy and associations with cognitive impairment in Parkinson disease. Neurology. 2017;88(13):1265–72.

35. Rash J, Yasumura T, Davidson K, Furman C, Dudek F, Nagy J. Identification of cells expressing Cx43, Cx30, Cx26, Cx32 and Cx36 in gap junctions of rat brain and spinal cord. Cell communication & adhesion. 2001;8(4-6):315–20.

36. Schwab BC, Heida T, Zhao Y, van Gils SA, van Wezel RJ. Pallidal gap junctions-triggers of synchrony in Parkinson’s disease? Movement disorders. 2014;29(12):1486–94.

37. Schwab BC, van Wezel RJ, van Gils SA. Sparse pallidal connections shape synchrony in a network model of the basal ganglia. European journal of neuroscience. 2017;45(8):1000–12.

38. Kelley R, Flouty O, Emmons EB, Kim Y, Kingyon J, Wessel JR, et al. A human prefrontal-subthalamic circuit for cognitive control. Brain. 2018;141(1):205–16.

